# Caribbean golden orbweaving spiders maintain gene flow with North America

**DOI:** 10.1101/454181

**Authors:** Klemen Čandek, Ingi Agnarsson, Greta J. Binford, Matjaž Kuntner

## Abstract

The Caribbean archipelago offers one of the best natural arenas for testing biogeographic hypotheses. The intermediate dispersal model of biogeography (IDM) predicts variation in species richness among lineages on islands to relate to their dispersal potential. To test this model, one would need background knowledge of dispersal potential of lineages, which has been problematic as evidenced by our prior biogeographic work on the Caribbean tetragnathid spiders. In order to investigate the biogeographic imprint of an excellent disperser, we study the American *Trichonephila*, a nephilid genus that contains globally distributed species known to overcome long, overwater distances. Our results reveal that the American *T. clavipes* shows a phylogenetic and population genetic structure consistent with a single species over the Caribbean, but not over the entire Americas. Haplotype network suggests that populations maintain lively gene flow between the Caribbean and North America. Combined with prior evidence from spider genera of different dispersal ability, these patterns coming from an excellent disperser (*Trichonephila*) that is species poor and of a relatively homogenous genetic structure, support the IDM predictions.

## Introduction

Among the archipelagos, the Caribbean offers one of the best researched natural arenas for addressing biogeographic hypotheses (Ricklefs and Bermingham 2008). Caribbean islands are numerous and are of sufficiently varied ages and sizes to provide an historical context that generated interesting biogeographic histories of the organisms that inhabit them. An emerging issue that is relevant to organismal biology, lineage diversification, as well as biogeographic histories and patterns, is the degree to which variation in dispersal propensity can predict species richness of lineages (Laube et al. 2013; Borda-de-Água et al. 2017). The Caribbean is an ideal archipelago to pose these questions.

The recently reformulated Intermediate Dispersal Model of biogeography (hereforth IDM) (Claramunt et al. 2012; Agnarsson et al. 2014) posits that differences among comparable lineages in dispersal potential over long distances affects their levels of gene flow over discrete units, such as islands, and that this variation is reflected in species richness patterns among these lineages. For example, if lineages contain poor dispersers, these organisms rarely colonize remote islands, leading to overall low species richness. Conversely, those lineages that are biologically capable of long distance travel may maintain such a lively gene flow among islands, or between island and continent, as to severely restrict speciation. Finally, those organisms with intermediate dispersal potential get to be carried to remote island rarely enough so that their founding populations may start to speciate, a hypothetical scenario that may result in the highest species richness. What this model implies is that biological attributes that define higher taxa, say genera, may link to the overall potential how these organisms disperse, and therefore affect their species richness, and biogeography.

The IDM, therefore, predicts species variation in richness among lineages to be a consequence of varying dispersal potential. However, in order to test the general validity of the model, one needs to identify appropriate test lineages. Ideally, these would be co-distributed in an archipelago, be of comparable taxonomic ranks, and furthermore exhibit a measurable variation in phenotypes that pertain to dispersal. This has rarely been done, as studies testing the IDM have mainly focused on either only excellent dispersers (Claramunt et al. 2012) or solely poor dispersers (Pabijan et al. 2012), on lab reared organisms (Venail et al. 2008), or they included multiple lineages of incomparable taxonomic ranks (Agnarsson et al. 2014). In this vein, our prior work has compared a tetragnathid spider lineage with a hypothetical low dispersal potential (*Cyrtognatha*) (Čandek et al. 2018a) with its close relative (*Tetragnatha*) (Čandek et al. 2018b) over the Caribbean and the mainland. We found *Tetragnatha* to be extremely species rich in the Caribbean and attributed this richness to a biology that has elements of excellent dispersal, mixed with repeated secondary loss of dispersal ability, all this resulting in a mixed pattern of cosmopolitan, as well as narrowly endemic lineages. In comparison, *Cyrtognatha* was relatively species poor, with exclusively single island endemic species. We here explore another lineage that we *a priori* expect to show excellent dispersal ability that is co-distributed over the Caribbean.

Our present study thus focuses on the nephilid genus *Trichonephila* Dahl in the Caribbean. *Trichonephila* is a global genus of golden orbweavers (Nephilidae) that contains species known to readily cross long, overwater distances (Kuntner et al. 2018). Only two species are known in the New World, with *T. clavipes* distributed widely from North to South America (Kuntner 2017). We aim to test this single species status in Americas through a Caribbean transect, and with numerous terminals from other parts of the New World. We predict that in accordance with the IDM, *Trichonephila* will be species poor compared with the above tetragnathids, and will show the least structured genetic pattern over the archipelago. If so, this would implicate a lively gene flow over all the islands.

## Results and Discussion

Our analyses reveal a phylogenetic and population genetic structure of *T. clavipes* consistent with a single species over the Caribbean. While *T. clavipes* is monophyletic, species delimitation analyses detect more than a single species over the Americas. Using the best fit model, GMYC delineates two species (marked as A and B in Fig. 1). The vertical bar in Figure 1 indicates the threshold where GMYC model shifts from phylogenetic Yule processes to coalescent population processes. Species A contains a dichotomy with a subclade containing the Caribbean and mainland North American populations, and a subclade with Colombian and Costa Rican populations. Species B is native to South and Central America. While species delimitations using some GMYC and a ABGD agree with the two species, alternatives exist (see Supplementary Figs. S1-S14). More precisely, some GMYC, mPTP and some ABGD suggest more than two species. However, these alternatives are less credible because they also suggest unrealistic species lumps of morphologically well-diagnosed species on other continents, such as *T. komaci* and *T. sumptuosa* being detected as a single species. Our conclusion that *T. clavipes* may in fact contain more than a single, Panamerican species, contradicts the classical morphological taxonomy (Kuntner 2017) but is consistent with a population genetic study in South America (Bartoleti et al. 2018).

**Figure 1:**
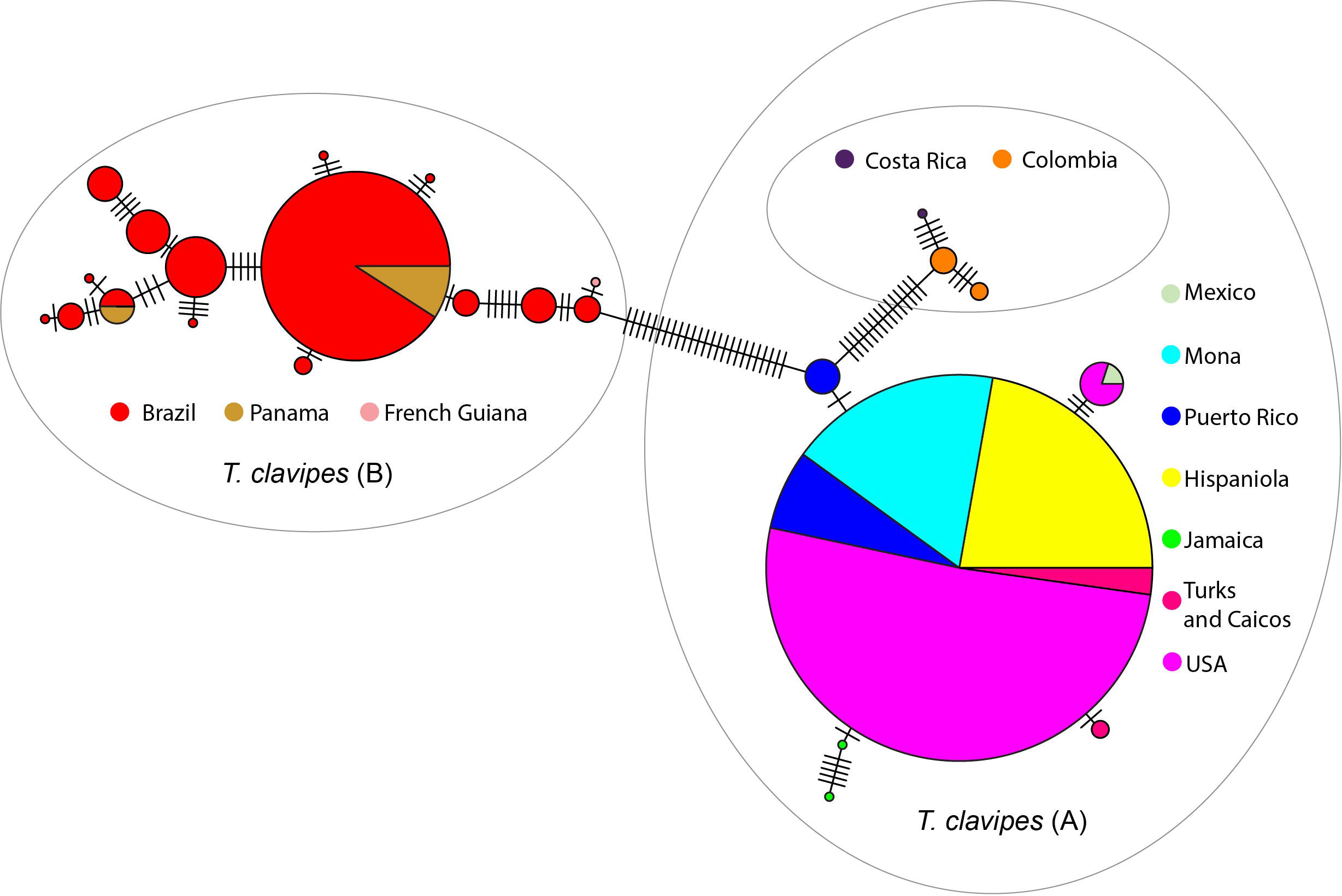
GMYC species delimitation splits *T. clavipes* into two species. Species (A) shows pronounced population structure with Caribbean + North America subclade and Colombia + Costa Rica subclade. Species B is South and Central American. Vertical bar represents a likely threshold for speciation processes in *Trichonephila*.

The haplotype network (Fig. 2) depicts a single, well-represented haplotype present on the sampled Caribbean islands, as well as in continental North America. A few point mutations separate this highly frequent haplotype with those present on Jamaica, Mexico, Puerto Rico, and Turks and Caicos. Other haplotypes are more distant, and form two distinct groups, one in Colombia and Costa Rica (putatively conspecific with the Caribbean), and another in Brazil, Panama and French Guiana that corresponds to the above species B. This haplotype network is consistent with two or three species of *T. clavipes*. It also strongly suggests that the Caribbean populations maintain gene flow with the North American mainland.

**Figure 2:**
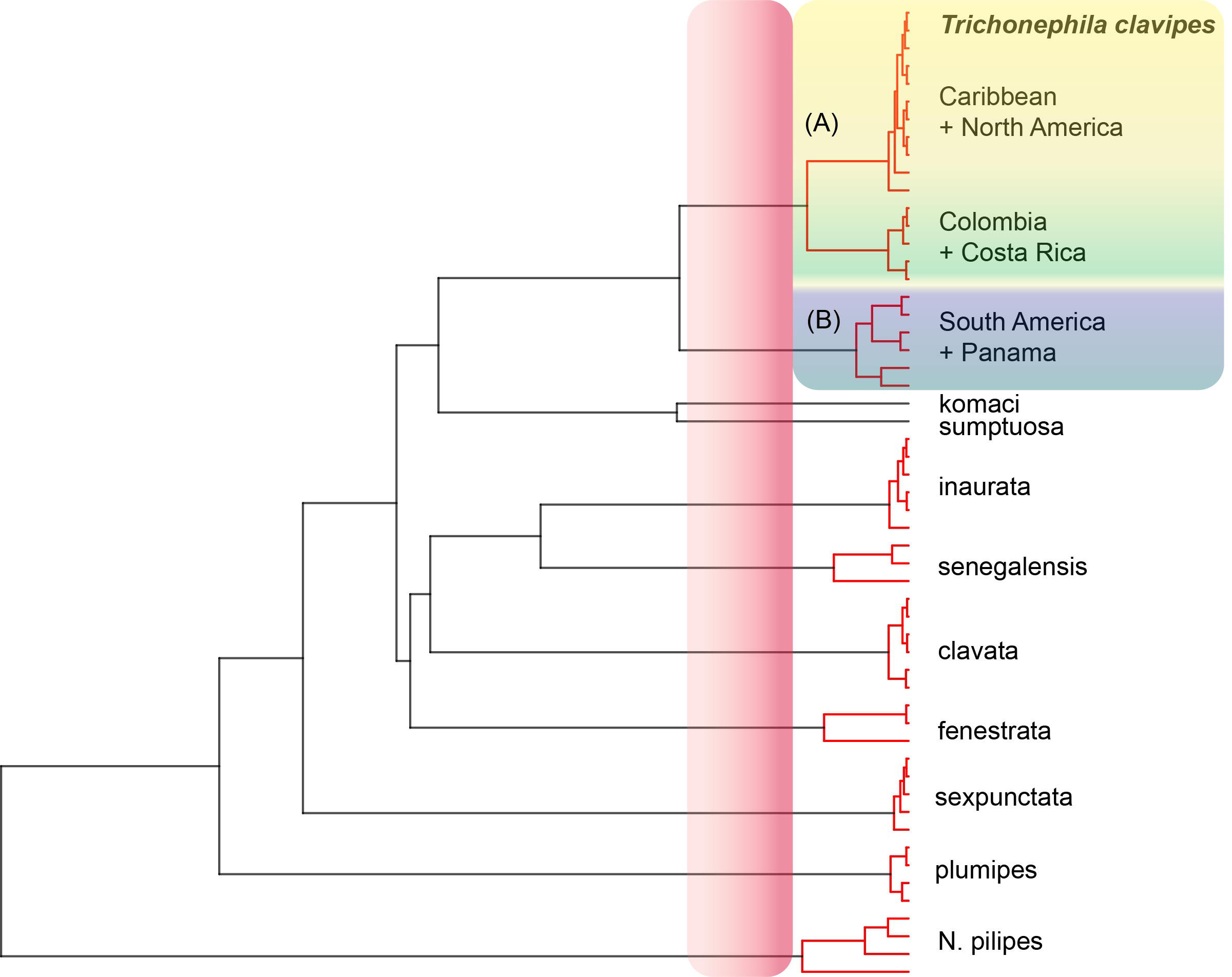
Haplotype network hints at two, or three species, of *T. clavipes*. Specimens from the Caribbean and North America form one well represented haplotype. Other, minor haplotypes are separated by very few mutations, suggesting high levels of gene flow in the area of interest.

These data reinforce the known biology of *Trichonephila* species as excellent dispersers. Although ballooning has not been directly observed in *Trichonephila*, it has been shown in the closely related and similar *Nephila* (Lee et al. 2015). Ballooning is an effective means of overcoming oceanic and continental barriers to gene flow, as several studies on Asian *Nephila pilipes* show (Lee et al. 2004, 2015; Su et al. 2006). More close relatives of *T. clavipes* also show extremely wide, genetically poorly structured population patterns worldwide. Good examples are *T. inaurata* that maintains gene flow between Africa and the islands of the Western Indian Ocean (Kuntner and Agnarsson 2011), as well as *T. edulis and T. plumipes* reported to travel seasonally from Australia to New Zealand (Harvey et al. 2007). There also seems to be a constant gene flow between the Korean and the Japanese populations of *T. clavata* (Jung et al. 2006).

*Trichonephila clavipes* resembles the Caribbean pattern detected in the araneid *Argiope argentata* where island populations clearly interbreed (Agnarsson et al. 2016). At a higher taxonomic level and within the area of interest, the Caribbean *Trichonephila* contrasts the two tetragnathid lineages: *Cyrtognatha* is a relatively poor to intermediate disperser with significant species richness and high endemism, *Tetragnatha* is a dynamic disperser lineage, with species apparently ranging from extremely good to relatively poor, and shows a high species richness and a mixed endemic to widespread mix of species. *Trichonephila* is an excellent disperser with a single species over the archipelago, exhibiting little genetic structure. Although *Tetragnatha* is an unusual case with apparently frequent evolutionary chance in dispersal potential, this triplet of genera provides preliminary support of the IDM.

In conclusion, *T. clavipes* does not seem to represent a single species over the Americas. The nominal *T. clavipes* is from the Caribbean (type is from Jamaica), but this species also inhabits North America, as well as parts of Central and South America. Over this vast geographical area, *T. clavipes* populations maintain a lively gene flow, suggesting that these spiders undertake airborne travel over the archipelago.

## Methods

### Dataset assembly

Our total dataset contains every available *T. clavipes* cytochrome c oxidase subunit 1 (COI) sequence from the combined Caribbean + USA region (N = 58), an equal number of COI sequences randomly selected from Brazilian *T. clavipes* (Bartoleti et al. 2018), and every available sequence of *T. clavipes* from other areas (4 x Panama, 4 x Colombia, 1 x French Guiana, 1 x Costa Rica, 1 x Mexico). We also targeted other *Trichonephila* global exemplars (1 to 6 terminals per species, 8 species total), and four individuals of *Nephila pilipes* as the outgroup (Supplementary table S1). We emphasized the richness in geographic terminal coverage over that of using more genes with fewer terminals. Using COI marker to resolve phylogeographic questions is a common, and valid approach (Su et al. 2006; Čandek and Kuntner 2015; Bartoleti et al. 2018).

### Phylogenetic reconstructions

To obtain ultrametric phylogenies, we performed Stepping Stone sampling and Bayes Factor test (Baele et al. 2012) within BEAST2 (Bouckaert et al. 2014). We reduced the total dataset (Supplementary table S1) and constrained the topology according to a phylogenomic hypothesis (Kuntner et al. 2018). Model tests (Supplementary note S2) selected a strict clock with the rate 0.0112 (Bidegaray-Batista and Arnedo 2011) and a coalescent constant population tree prior. We used bModelTest (Bouckaert and Drummond 2017) as nucleotide substitution model.

We reconstructed a Bayesian phylogeny using the reduced dataset as above in MrBayes (Huelsenbeck and Ronquist 2001), running four independent MCMC chains with 30 million generations, 25 % burn-in and a sampling frequency of every 1000. GTR+G+I was the nucleotide substitution model (Darriba et al. 2012). Finally, we ran another Bayesian phylogeny for the *T. clavipes* ingroup only, with settings as above, except with 10 million generations.

### Species delimitations

For species delimitation analyses, we employed three methods, the Generalized Mixed Yule Coalescent (GMYC) (Fujisawa and Barraclough 2013), the Multi-rate Poisson Tree Processes (mPTP) (Kapli et al. 2017) and the Automatic Barcode Gap Discovery (ABGD) (Puillandre et al. 2012). We ran GMYC delimitations in “splits” package of R version 3.5.1. (R Core Team 2018), using the utrametric tree and testing single, as well as multiple thresholds settings. We ran mPTP delimitations online using default settings and with ultrametric as well as Bayesian trees. For ABGD delimitations, we uploaded fasta sequences to its online platform and tested all three implemented substitution models (JC69, K80 and Simple distance). Here, we present a GMYC delimitation result using the best model for the data (Fig. 1) while 14 additional delimitation results are in the supplementary material (Supplementary Figs. S1 to S14).

### Haplotype network reconstruction

We used “pegas” package in R to reconstruct a haplotype network of all 127 *T. clavipes* individuals from 12 areas. We trimmed the sequences to equal lengths, resulting in 537 remaining nucleotides. The sizes of circles in the reconstructed network correspond to the frequency of a specific haplotype.

## Data accessibility

All data and protocols needed to replicate this study are included in the paper and its Supplementary Material.

## Contributions

Research design, material acquisition, molecular procedures: All authors. Data analyses: K.Č.K.Č. and M.K. wrote the first draft of the paper. All authors contributed to writing and revising the paper.

## Acknowledgements

We thank the entire CarBio team (http://www.islandbiogeography.org/participants.html) for collecting the material across the Caribbean. Moreover, we thank Lisa Chamberland and other members of the Agnarsson lab (http://www.theridiidae.com/lab-members.html) for the help with material sorting and molecular procedures.

